# Viral miRNAs Confer Survival in Host Cells by Targeting Apoptosis Related Host Genes

**DOI:** 10.1101/2020.06.10.144469

**Authors:** Md. Sajedul Islam, Abul BMMK Islam

## Abstract

**Background:** miRNAs are small non-coding RNAs that regulate the expression of genes by RNA silencing method. Like eukaryotic organisms, some viruses also produce miRNAs. While contribution of host miRNA in the prevention of viral pathogenesis has been studied, it is not known very well how viral miRNA can confer its survival in the host. Here we hypothesized that viral miRNAs can bind to the host target genes to confer their pathogenicity by down-regulating specific pathways and related genes that otherwise pose threat to cell survival.

**Methods and Results:** Using targets of 168 viral miRNAs from 13 different viruses overrepresentation analysis was done. Functional enrichment analysis of the genes targeted by the miRNAs indicates that viruses target specific immune system and host defense related pathways via miRNA mediated gene silencing. Integration and analysis of the publicly available experimental host gene expression data by RNA-seq provided insight that viruses target host apoptosis process by switching off related genes through miRNA induced mechanisms and thus probably ensure their survival.

**Conclusions:** As switching off the apoptosis of host cells would provide the viruses with selective advantages in surviving inside host, our findings therefore envisage an important function of viral miRNA which demands further *in vivo* experiments for better understanding in this regard.

## 1. Introduction

miRNAs are ∼22 nucleotide, small, non-coding RNAs that are present in the vertebrates, plants, invertebrates and in a wide range of viruses [1]. The primary function of miRNAs is to regulate the expression of genes post-transcriptionally, via base pair formation with the 3’-Untranslated Region (3’-UTR) of specific messenger RNAs (mRNA). Viral miRNAs play subtle roles in the survival and proliferation of viral particles through host immune system evasion [2], establishing microenvironment for viral replication [3], regulation of innate immune system, differentiation of adaptive immune cells [4] etc.

Like most animals and plants, some viruses (mostly DNA viruses), can also produce miRNAs that provide them different selective advantages required for their survival, host immune system evasion, regulation of host and viral genes [5], viral replication [6], influencing viral latency [4], diminishing host antiviral responses, suppressing host cell apoptosis [5] etc. Initially, viral miRNAs were found in the Epstein-Barr virus [7] but later on other viral families like, Herpesvirus, Polyomavirus, Ascovirus, Iridovirus, Baculovirus, Adenovirus, Retrovirus [8] etc. were also found to produce miRNAs [9]. At present more than 20 human viruses are expected to produce miRNAs, among which almost 10-12 are already validated experimentally. These human viruses produce almost about 135 miRNAs [10], which are thought to be beneficial for the viruses to survive in human bodies.

These viruses produce miRNAs that can control the expression of the corresponding viral genes and some human genes as well. The miRNA sequences found from these viruses have been analyzed and their respective target sites have been found at the 3’-UTR regions of many genes in human [10]. It is expected that these miRNAs may provide some selective advantages to the viruses so that they can evade the host defense machineries. Since viruses need to replicate inside host cell, they may need to take necessary initiative to prevent host cell death by apoptosis and other immunologic mechanisms. That is why, we hypothesized that viral miRNAs may down regulate apoptosis by targeting apoptosis related genes, thereby they ensure prolonged survival of the host cells as well as themselves.

## 2. Materials and Methods

### 2.1. Identification of Viral miRNAs and their Target Genes

To understand the functions of viral miRNAs in providing selective advantages to the viruses, first it was required to know the viruses which produce the miRNAs. For this purpose miRbase [11] was explored and the human viruses that produce miRNAs were identified. From miRBase mature FASTA sequences of the viral miRNAs were also extracted for further analyses. Using the target identifying tools TargetScan [12], RNAhybrid [13] and PITA [14] we scanned the 3′-UTR regions of the human protein coding genes for putative binding of miRNAs and from databases UTRdb [15], miRTarBase [16] that house target genes of viral miRNAs we obtained the putative and experimentally validated target gene sequences, gene symbols etc. Target genes obtained from these databases and targets that we identify by scanning UTRs were combined to make a unique union set of viral miRNA targets.

### 2.2. Functional Enrichment Analysis

To annotate functions of all the genes we obtained the Biological Processes (GOBP), Cellular Locations (GOCL) and Molecular Functions (GOMF) data from the Gene Ontology Consortium (GO) [17, 18]. The pathways involved were obtained from Kyoto Encyclopedia of Genes and Genomes (KEGG) [19]. In both cases Ensembl genes (Release 79: March, 2015) [20] were used.

We used Gitools [21] (Version: 1.8.4) to perform enrichment analysis and to generate heatmaps using the targeted biological processes or pathways and the corresponding p-values. The Right P-value was used as the parameter. Genes associated with the biological processes and pathways can be viewed and thereby the target genes in each biological process or pathway can be observed.

### 2.3. Gene Expression Microarray Data Analysis

Gene Expression Omnibus (GEO) [22] is a curated, public reservoir of microarray gene expression data at National Center for Biological Information (NCBI). From GEO, we used the RNA-seq dataset GSE44769 which used the platform Illumina Genome Analyzer IIx (GPL10999). In this particular dataset, the experiment was done using Epstein-Barr virus (EBV) cell line in four different conditions including “No BART, EtOH”, “BART, Dox”, “No BART, Dox” and “Dox, EtOH” to obtain the expression of genes in those conditions. The data of “No BART, EtOH” was taken as the control of this dataset and the data of “BART, Dox” was taken as the most appropriate sample. Using in house R script the data were converted into Log2 values from absolute expression and the differential expression level of the genes in the experimental condition were identified comparing with the control.

“Apoptosis” related gene set was extracted from KEGG database “apoptosis pathway” terms and the expressions of those genes were obtained from the RNA-seq data. Apoptotic genes which were down-regulated more than two folds were taken in consideration to identify viral miRNAs targeting them. We studied the functions of these significantly down-regulated genes to learn about the selective advantages that the miRNAs can provide the viruses by knocking down these genes. Overview of the complete research is summarized in **Figure-1**.

**Figure-1:**
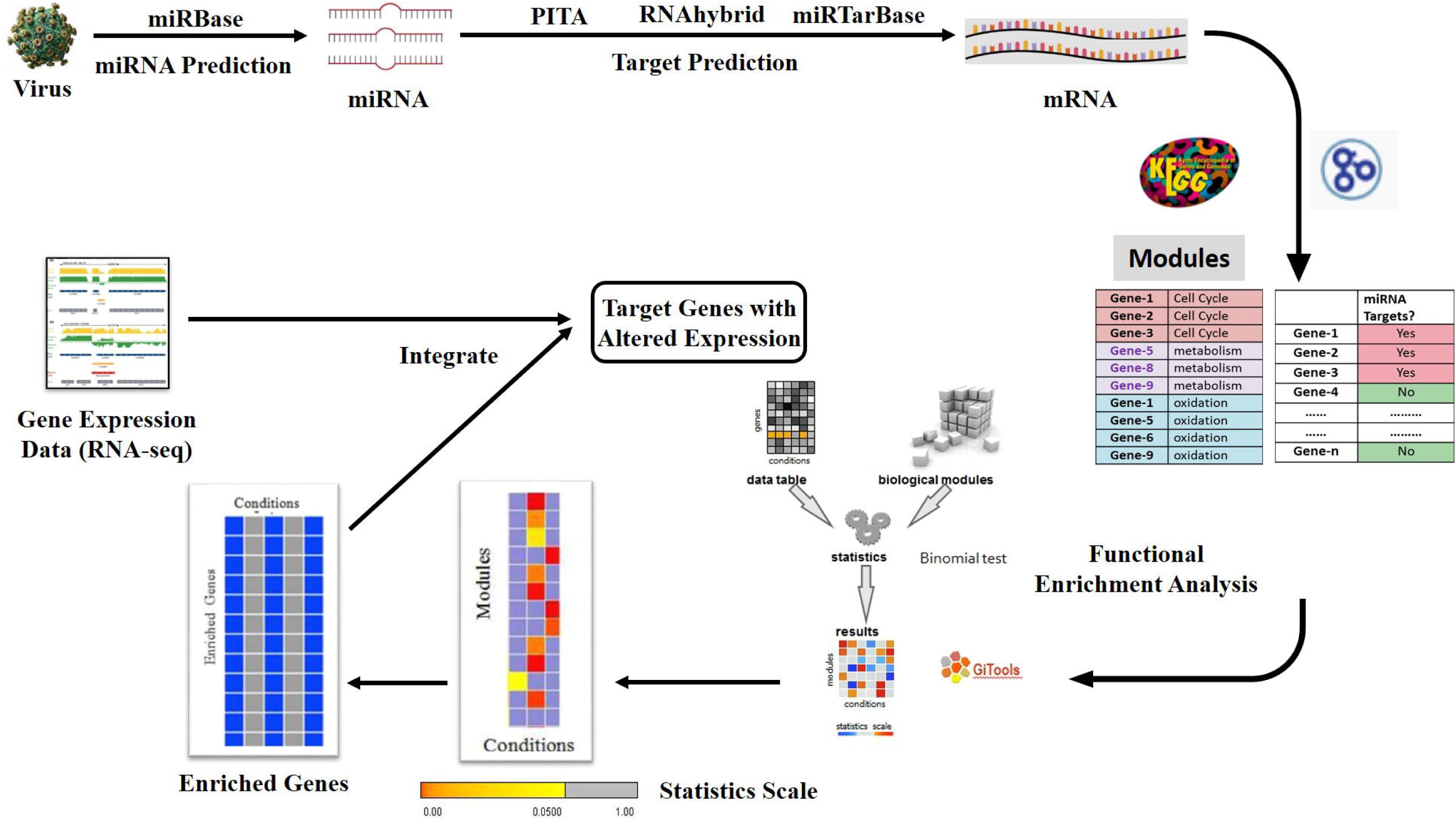
Schematic diagram summarizing the study.

## Result

### 3.1. From 13 Human viruses 168 miRNAs and their targets were retrieved from various databases

From miRbase and various publications [4, 5, 7, 10, 23-27], miRNAs of the human viruses were obtained. Among these viruses, 13 human viruses were chosen whose putative miRNAs were already published (**Additional File-1**). There were 168 miRNAs observed from these 13 viruses. Taking these viral miRNAs with their sequences obtained from miRBase, target genes in human genome were identified from the databases RNAhybrid [13], miRTarBase [16] and from the online software PITA [14] and TargetScan [12]. A union set of target genes was created combining all the obtained target genes from the databases and tools. There were almost 22,000 unique genes observed which were targeted by the viral miRNAs, among which, EBV targeted about 16,000 genes, HIV, HBV, HSV1, HSV2, KSHV and HCMV targeted about 15,000 genes, BKV, JCV and MCV targeted about 7,000 genes.

### 3.2. Functional enrichment analysis shows clusters of over-expressed pathways and biological processes

Enrichment analysis is performed to evaluate the statistical significance of biological conditions and to identify whether particular sets of biological processes are targeted in an increased proportion. The biological processes and KEGG pathways which were significantly enriched were taken into consideration for preparing different clusters. The cut-off value for the significance was set to FDR corrected P-value 0.05 (**Figure-2**).

**Figure-2:**
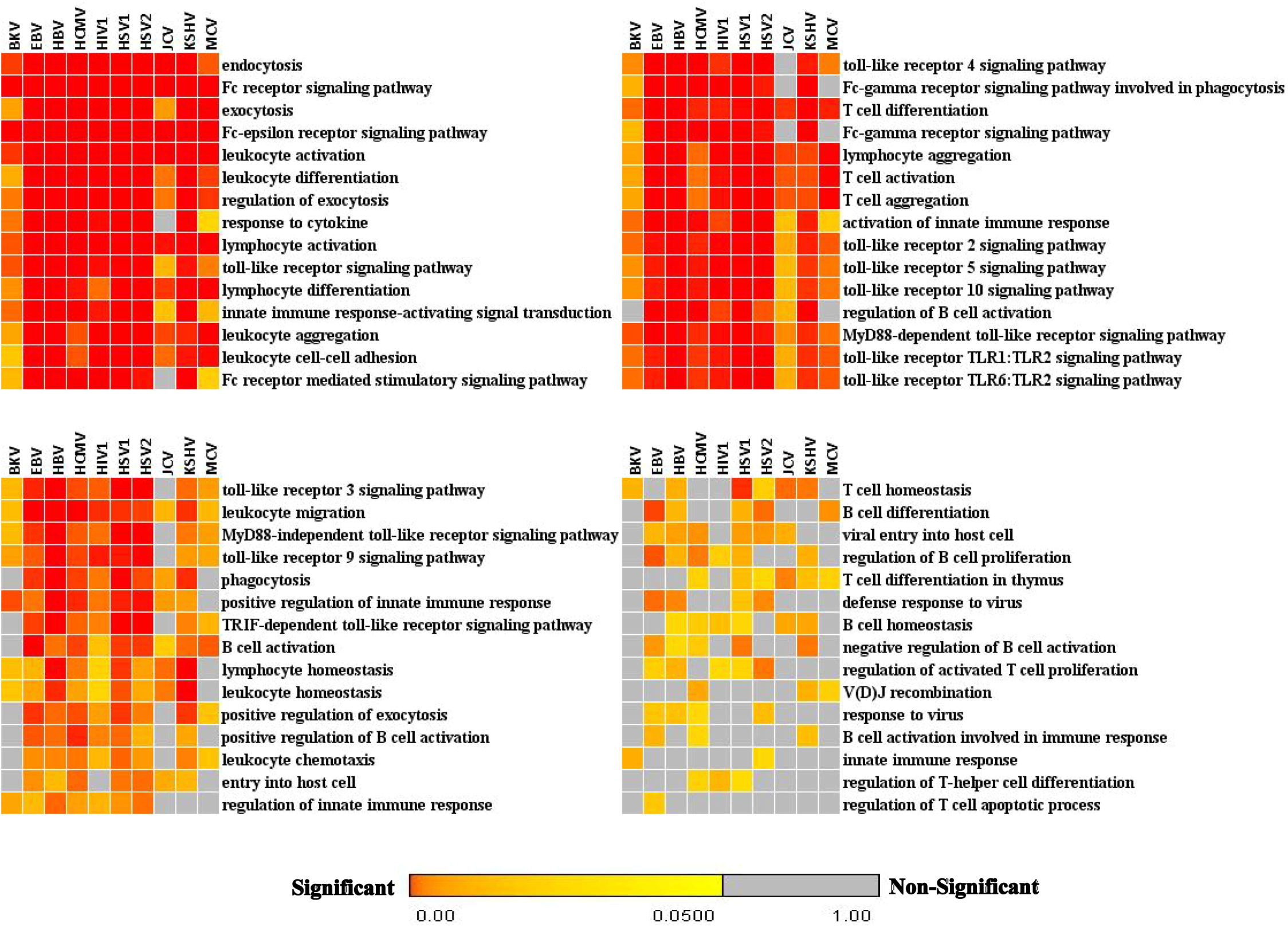
Enrichment of gene ontology biological processes: Representation of (selected) GOBP (Immune system) enriched terms related to immune system obtained from Gitools. Statistical significance (FDR corrected p-values) is represented in a color coded scale. Color towards red indicates more significant and color towards yellow means less significant, and gray color indicates non-significance (> 0.05).

By studying various experiments and related articles several important KEGG pathways and associated biological processes were identified which were clustered together and selected for further analyses. The selected pathways were: Calcium mediated signaling, MAPK cascade, Wnt signaling pathway and Apoptosis (**Figure-3**).

**Figure-3:**
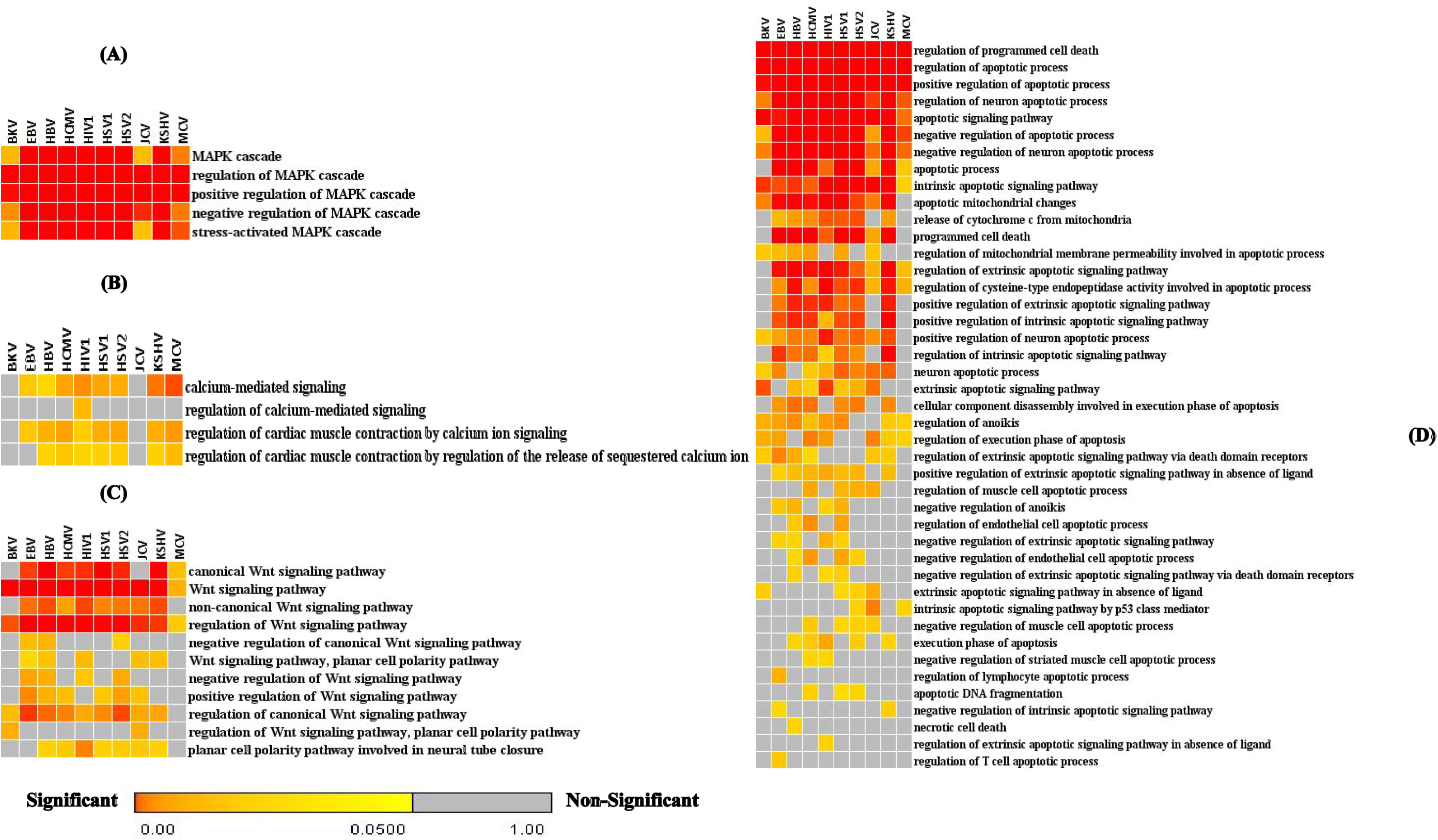
Enrichment analysis of pathways: Overrepresentation of Biological processes of (A) MAPK cascade, (B) Wnt signaling pathway, (C) calcium mediated signaling, (D) Apoptosis obtained from Gitools. Color scale and legend as in Figure 2.

### 3.3. Enrichment analysis of Apoptosis provided 84 enriched genes targeted by viral miRNAs

Enrichment analysis provided us the enriched biological processes related to apoptosis. We then identified the annotated enriched genes of the apoptosis process from KEGG pathway in Gitools. There were 84 enriched genes (corrected p-value < 0.05) observed which were targeted by the viral miRNAs. Enriched apoptosis related genes were retrieved and represented in a color coded heatmap (**Figure-4**).

**Figure-4:**
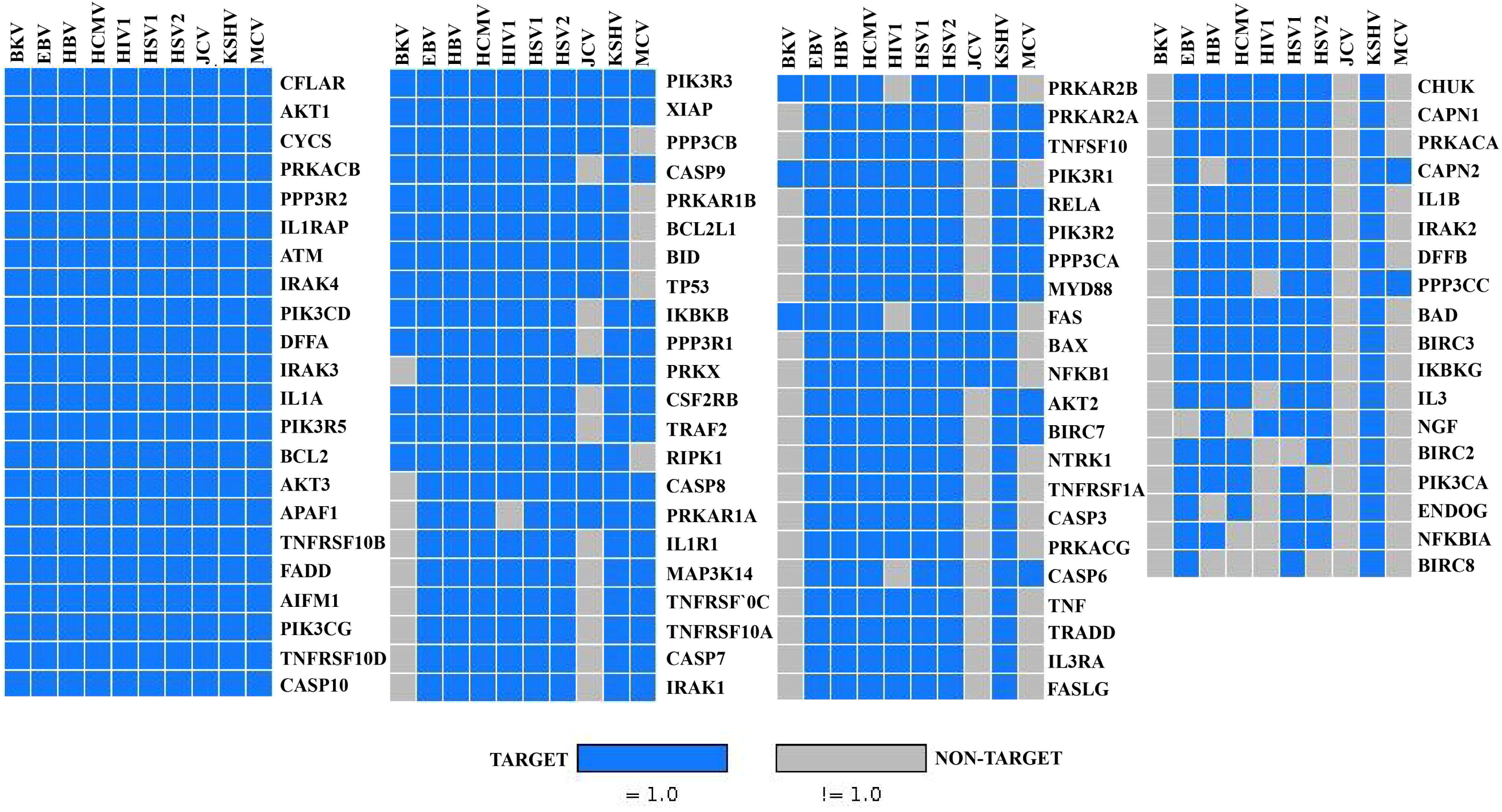
Enriched apoptosis genes targeted by viral miRNAs: Enriched genes related to apoptosis process obtained from Gitools. The Blue colored hits express the genes that are targeted by viral miRNAs and the Gray colored hits are the genes that are not targeted by viral miRNAs. Vertical lines express viral miRNAs and the horizontal lines express corresponding target genes.

### 3.4. Microarray analysis of EBV cell line provided significant downregulation of 49 genes associated with Apoptosis

Analyzing the Microarray data (GSE44769) obtained from GEO we found that 75 apoptosis genes that showed deviated expression level compared to the control condition. Among these 75 genes, 24 genes were up regulated and the rest 51 genes were down regulated. As we assumed that viral miRNA would down-regulate apoptosis process, we selected the genes which were down-regulated at least two-fold. We observed 49 such genes when they were compared to the control (**Figure-5**).

**Figure-5:**
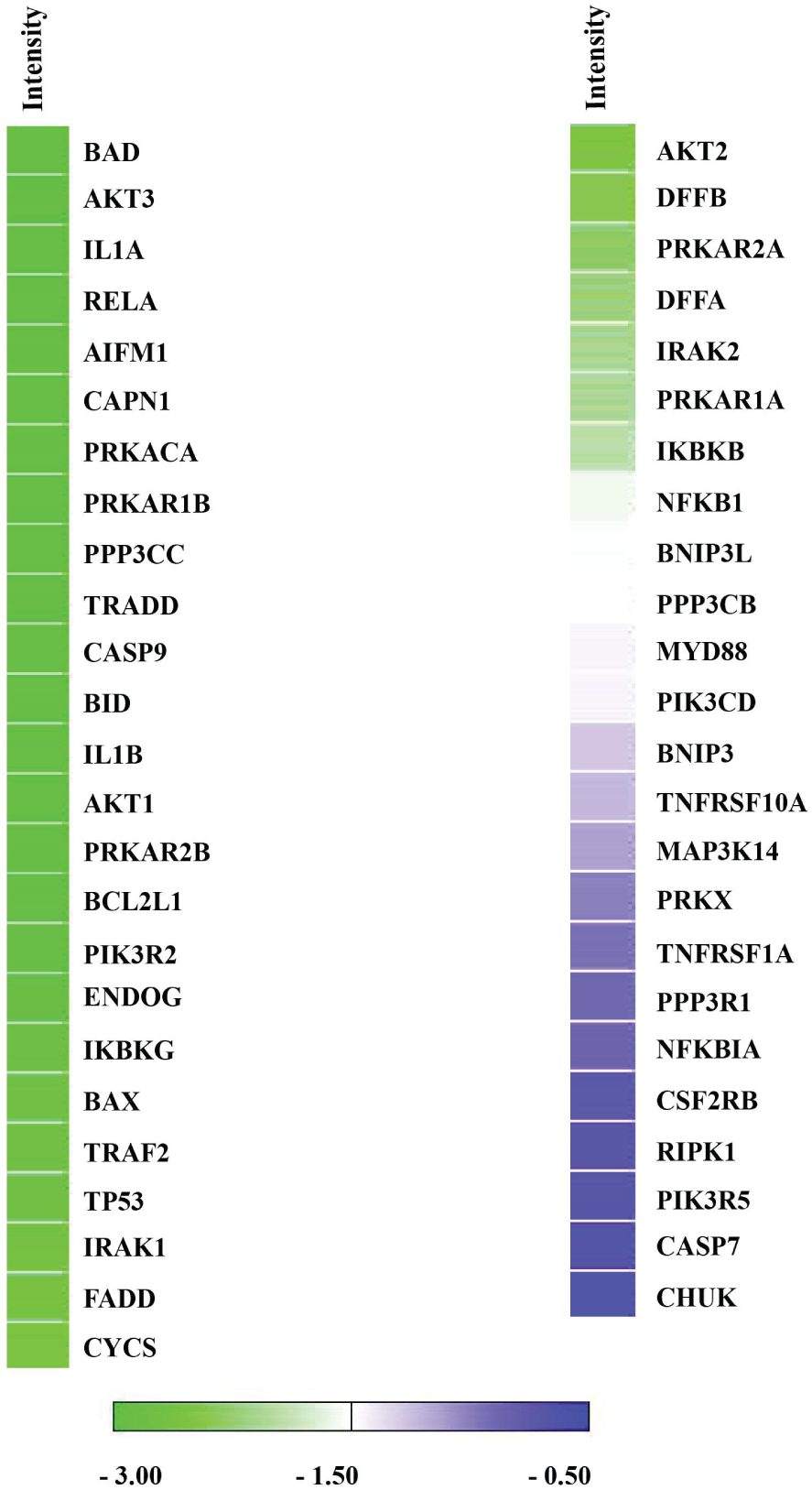
Expression level of the down-regulated genes: Down-regulated genes are represented in a color coded scale based on their expression level. The genes that are down-regulated at least two-fold (Log2 ratio < -0.5) are considered. The significance level of down-regulation is taken at -1.5 of the Log2 ratio, which is represented as White color. Genes having Log2 ratios less than -1.5 are represented as Green color and genes having values more than -1.5 are represented as Blue color.

After getting the down regulated genes from the RNA-seq we assigned the miRNAs of EBV to their corresponding target genes with their expression levels. We found out that there were several miRNAs which targeted same apoptosis gene (**Additional File-2**).

### 3.5. Functions of the 20 most significantly downregulated apoptosis genes provided insight about the reason of miRNAs targeting them

As soon as the EBV miRNAs were assigned with their corresponding apoptosis genes, the most significantly down regulated apoptosis genes were identified from the total genes in the Microarray data taking the bottom 20% expressed genes (Log2 ratio < -2.74). There were 20 apoptosis genes observed which were most significantly down regulated (**Additional File-2**). Functions of these genes were studied from UniProt [28, 29] protein database to learn about the reason of miRNAs targeting them (**Table-1**).

**Table-1:**
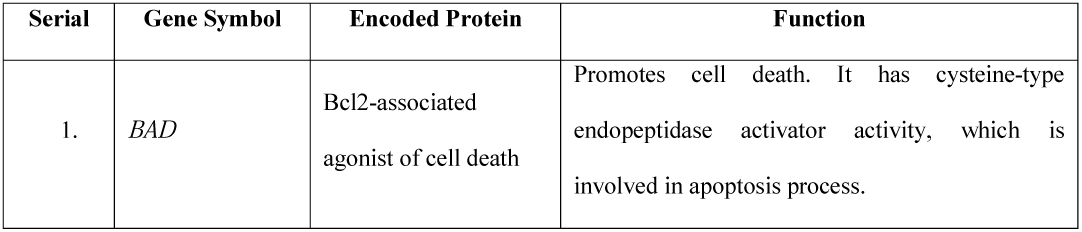
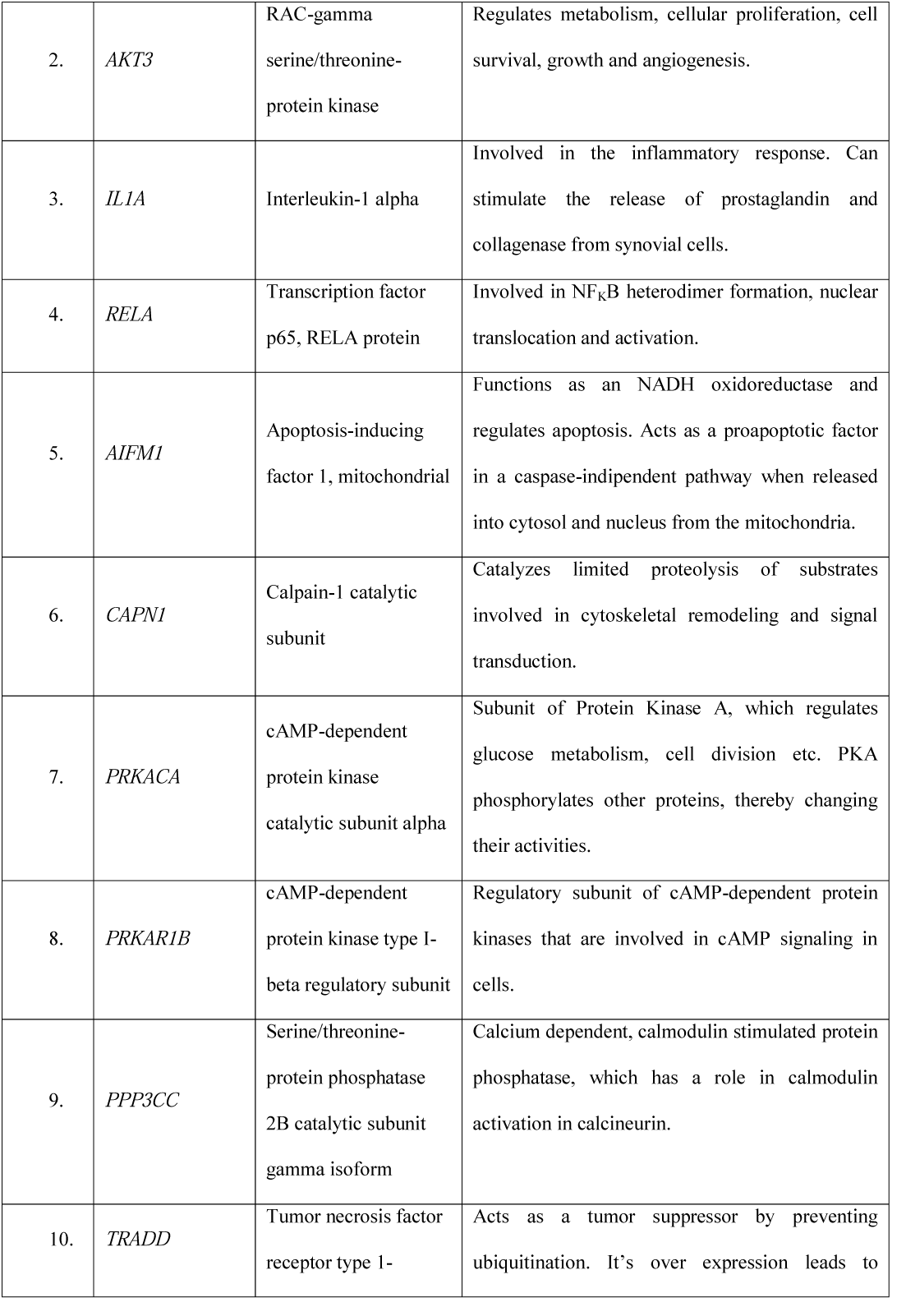
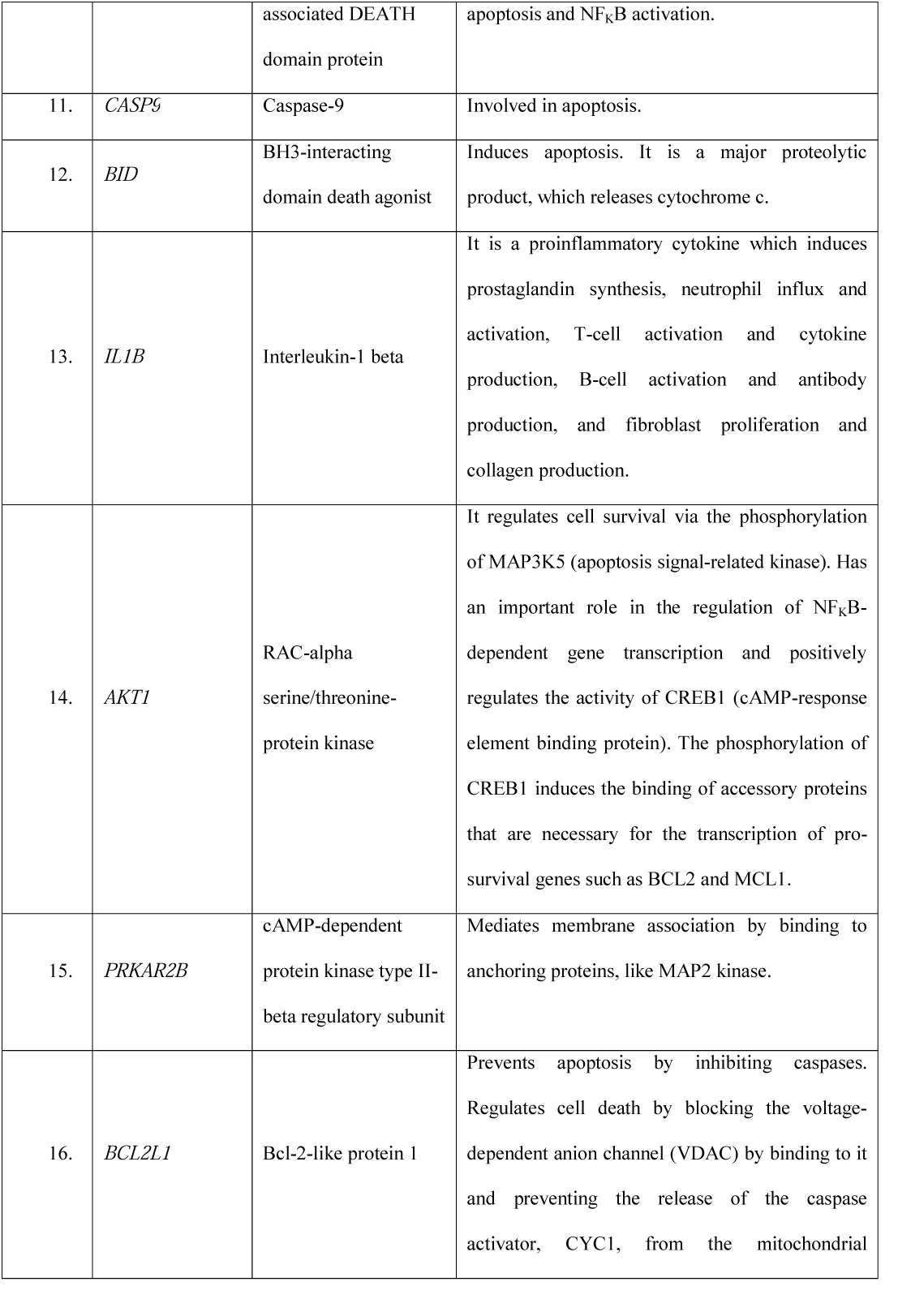
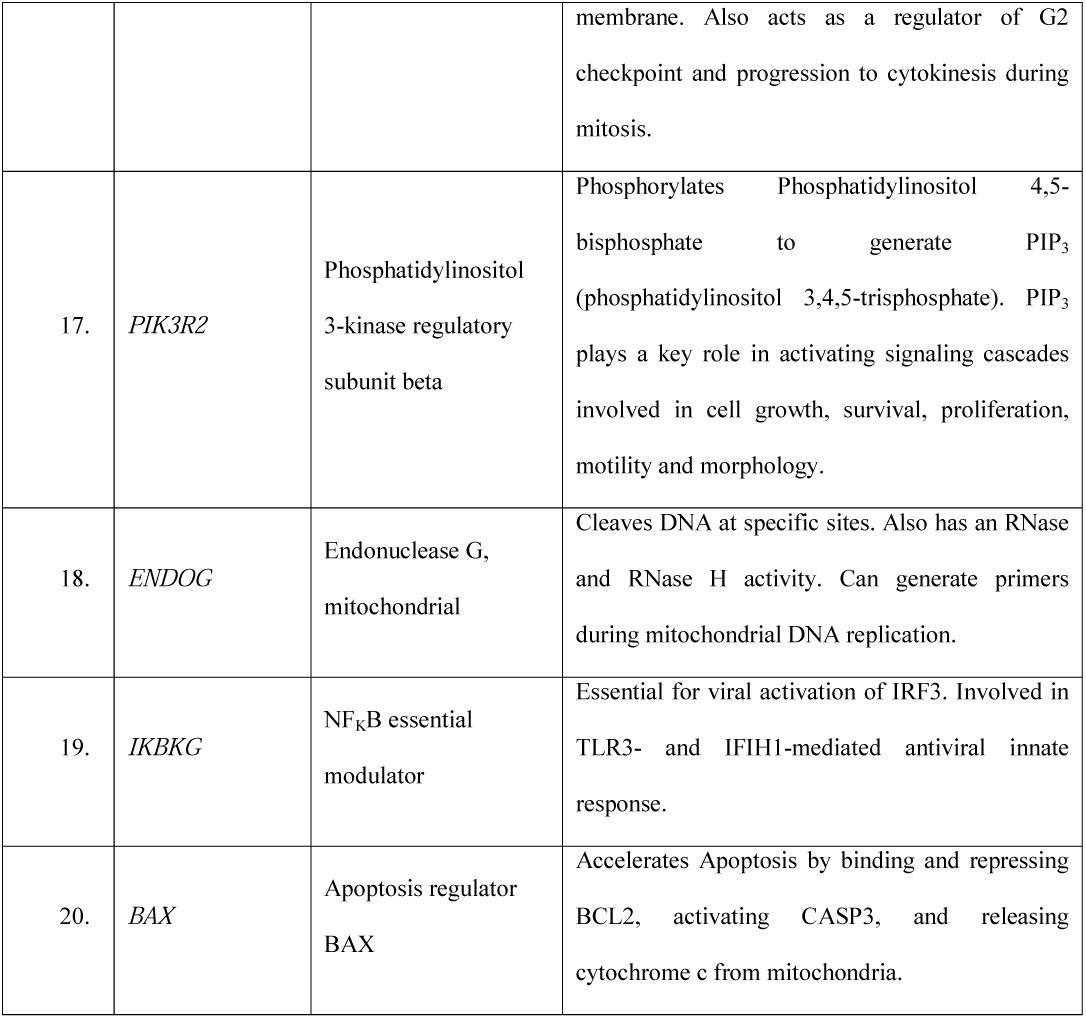
Functions of most significantly down regulated genes obtained from UniProt

## 3. Discussion

miRNAs have become an important subject of interest in recent years in the fields of molecular biology. Their roles in regulating gene expression have become something of remarkable importance to learn about the functions of particular genes and their encoded products. Various organisms including vertebrates, invertebrates [1], plants [1], viruses etc. produce miRNAs to regulate the expression of their genes, thereby controlling their physiological functions. At present there are more than 9000 miRNA genes annotated among which about 700 miRNA genes are from humans [30]. These miRNAs have various roles to ensure proper homeostatic conditions of an organism. One such role includes the host-pathogen relationship.

It is already established in various experimental works that human miRNAs target viral genes [31, 32] and function as antiviral mediators to suppress viral pathogenesis. By silencing the disease-causing genes of a virus human miRNAs ensure the prevention of any deleterious events caused by that virus. On the contrary there are very little evidences to prove that viral miRNAs effectively target and regulate host genes. Pathogenic viruses cause several diseases in human and human defense machineries continuously encounter and remove these pathogenic viruses from the system. To evade these host defense molecules viruses might have further evolved to produce miRNAs to silence host genes. This silencing can provide them with various selective advantages including host defense evasion, viral replication [5], diminishing antiviral responses etc. To accentuate this event whether viruses effectively target and control host genes we proceeded with several scientific works from different laboratories and gained insight about the role of viral miRNAs in their own survival.

After performing enrichment analysis, we observed that various physiological processes are enriched at a high level. We clustered these pathways and found viruses target biological processes like immune system related pathways, Wnt pathway, MAP Kinase cascade, apoptosis etc. at a significantly high level through their miRNAs. We predicted that switching these pathways off may serve them some selective advantage in their survival inside human host. So, we studied about these pathways and other related findings.

*Carl et al*. [10] worked with several viruses and associated miRNAs to hypothesize that viral miRNAs target host pathways by regulating the host gene expressions. They looked for the predicted miRNAs from different databases and tried to find the target genes. Then they obtained the enriched GO biological processes and predicted that viral miRNAs target some pathways. Their major drawback was that they failed to provide any experimentally validated proof in these regards. We identified the pathways and their corresponding genes targeted by viral miRNAs which were further verified by using RNA-seq datasets of *Vereide et al*. [33] who worked with miRNAs of Epstein-Barr virus and found that some EBV miRNAs are responsible for the transformation of B lymphocytes in Burkitt’s Lymphoma. We are providing here proof that viral miRNAs do target human genes and control specific pathways, like apoptosis.

Apoptosis is a natural physiological phenomenon where cells undergo programmed death when they reach maturity or are infected by any pathogen [34-36]. This process is very important to maintain homeostasis of an organism. Apoptosis prevents various deleterious effects like the progression of viral pathogenesis, cancer etc. As we have already found that viral miRNAs target host genes for their own survival, we predicted that viruses may specifically target and regulate host apoptosis process so that they can ensure their refuge and prolonged survival in host cells. Bearing this context in mind, we searched for the most significantly targeted genes by viral miRNAs and found that there are several host genes related to apoptosis which are down-regulated at a very significantly high level. These findings strengthened our hypothesis about viruses selectively targeting host apoptosis pathway for their own survival.

There are several viruses that cause various deleterious effects in humans. One such disease-causing virus is EBV, which is also called Human Herpesvirus-4 (HHV-4). EBV is the causative agent of a common disease named Infectious mononucleosis or simply Mono. But it can also cause several forms of cancer including Hodgkin’s lymphoma, Burkitt’s lymphoma, gastric cancer, nasopharyngeal carcinoma etc. There are several evidences showing that EBV is associated with some autoimmune diseases as well [37]. Despite causing so much adverse effects in human very less information is known about the complete pathogenesis of EBV inside host system. To gather insight about the pathogenesis of this particular virus we predicted that their pathogenesis may involve silencing the genes of the hosts via miRNAs.

*Pfeffer et al*. [38] published their scientific findings about the miRNAs of the viruses that belong to the Herpesvirus family. They performed computational methods to predict pre-miRNAs and then experimentally obtained the miRNAs produced by these viruses. We further worked with the miRNAs specially those of EBV and obtained their target genes. As mentioned earlier, we found several apoptosis related genes that are significantly controlled by EBV miRNAs. That accentuates the fact that one of the ways of viral pathogenesis includes miRNA mediated gene silencing. We studied the functions of the apoptosis related genes that are highly knocked down by viral miRNAs and found that most of them are associated with controlling apoptosis by positively influencing the process. Thus, it results in continued survival of the host cells and provides prolonged refuge to the viral particles when these genes are silenced by viral miRNAs.

Genes like *BAD, CASP9, BID* etc. are associated with the induction of apoptosis in human, which are highly targeted by viral miRNAs. Observing the experimentally validated expression levels of these genes after introduction of viral miRNAs we predict that viruses selectively target these genes that positively control apoptosis and switch them off by producing specific miRNAs. miRNAs are emerging to be important molecules in the fields of molecular biology, genetic engineering etc. It is gaining emphasis in the sectors of understanding the functions of genes and gene products. We can further experiment with these miRNAs to learn more about the host-pathogen interactions and combat the emerging pathogens. Diseases that are yet to be overcome due to lack of knowledge can be fought against with the knowledge of miRNA-gene interactions.

## 4. Conclusion

The role of miRNAs in regulating gene expression has become a significant aspect scientific interest recently. Here, we showed the role of miRNAs in the establishment of viruses inside the human hosts through controlling the apoptosis process and ensuring their prolonged refuge. This finding can be further strengthened by *in vivo* experiments that would result in better understanding in this context.

## List of Abbreviations

miRNA: microRNA;
UTR: Untranslated region;
DNA: Deoxyribonucleic acid;
GO: Gene Ontology;
BP: Biological processes;
CL: Cellular locations;
MF: Molecular functions;
KEGG: Kyoto Encyclopedia of Genes and Genomes;
GEO: Gene Expression Omnibus;
HIV: Human Immunodeficiency virus;
EBV: Epstein-Barr virus;
HBV: Hepatitis B virus;
HSV: Herpes Simplex virus;
KSHV: Kaposi’s Sarcoma-associated Herpesvirus;
HCMV: Human Cytomegalovirus;
BKV: BK Polyomavirus;
JCV: JC Polyomavirus;
MCV: Merkel Cell Polyomavirus;
FDR: False discovery rate.

## Acknowledgements

We acknowledge our gratitude for the research support by the Department of Genetic Engineering and Biotechnology, University of Dhaka.

## Availability of Data and Materials

The datasets supporting the conclusions of this article are included in the article and in the additional files.

## Funding

The project was not funded by any organization.

## Additional Files

**Additional File-1:**

- **Title of data:** Human viruses with miRNAs
- **File format:** Microsoft Word Document
- **Description of data:** Provides information about the human viruses that produce miRNAs

**Additional File-2:**

- **Title of data:** miRNAs of EBV and their target genes with expression levels
- **File format:** Microsoft Word Document
- **Description of data:** Provides information about the miRNAs produced by EBV and the genes targeted by them

## Notes

### Competing Interest Statement

The authors have declared no competing interest.

