## Additional File-1 for "Viral miRNAs Confer Survival in Host Cells by Targeting Apoptosis Related Host Genes"

**Human viruses with miRNAs**

| **Serial** | **Viruses** | **No. of mature miRNAs** |
| --- | --- | --- |
| 1 | BK Polyomavirus (BKV) | 02 |
| 2 | Epstein-Barr Virus (EBV) | 44 |
| 3 | Human Cytomegalovirus (HCMV) | 26 |
| 4 | Human Herpesvirus 6A (HHV 6A) | 01 |
| 5 | Human Herpesvirus 6B (HHV 6B) | 08 |
| 6 | Human Immunodeficiency Virus 1 (HIV 1) | 04 |
| 7 | Herpes Simplex Virus 1 (HSV 1) | 27 |
| 8 | Herpes Simplex Virus 2 (HSV 2) | 24 |
| 9 | Human Torque Teno Virus (TTV) | 01 |
| 10 | JC Polyomavirus (JCV) | 02 |
| 11 | Kaposi’s Sarcoma-associated Herpesvirus (KSHV) | 25 |
| 12 | Markel Cell Polyomavirus (MCV) | 02 |
| 13 | Simian Virus 40 (SV 40) | 02 |
