## Additional File-2 for "Viral miRNAs Confer Survival in Host Cells by Targeting Apoptosis Related Host Genes"

**miRNAs of EBV and their target genes with expression levels**

| **Serial** | **Gene Symbol** | **EBV miRNA** | **Expression Level** |
| --- | --- | --- | --- |
| 1. | BAD | BART13*, BHRF1-3 | -6.17948429 |
| 2. | AKT3 | BART10*, BART11-3p, BART13*, BART13, BART1-3p, BART14, BART15, BART1-5p, BART16, BART17-3p, BART17-5p, BART18-3p, BART19-3p, BART19-5p, BART20-3p, BART21-3p, BART22, BART2-3p, BART2-5p, BART3*, BART3, BART4*, BART4, BART5, BART6-3p, BART6-5p, BART7, BART8*, BART9*, BART9, BHRF1-2*, BHRF1-3 | -4.893248986 |
| 3. | IL1A | BART13*, BART15, BART1-5p, BART19-3p, BART20-3p, BART22, BART8*, BART9 | -4.496059485 |
| 4. | RELA | BART14*, BART14, BART17-3p, BART4, BART6-3p | -3.831753603 |
| 5. | AIFM1 | BART10, BART12, BART13*, BART13, BART14, BART17-5p, BART18-5p, BART2-5p, BART3, BART4, BART8* | -3.767094957 |
| 6. | CAPN1 | BART11-5p, BART12, BART13*, BART20-3p, BART2-3p, BART3*, BART6-3p | -3.530046006 |
| 7. | PRKACA | BART12, BART14, BART17-3p, BART17-5p, BART20-3p, BART2-3p, BART3, BART5, BART6-3p, BART6-5p | -3.484499608 |
| 8. | PRKAR1B | BART13*, BART13, BART17-3p, BART2-5p, BART4*, BART4, BART5, BART6-3p | -3.452551955 |
| 9. | PPP3CC | BART10*, BART14, BART8* | -3.450343372 |
| 10. | TRADD | BART4, BART5, BHRF1-2* | -3.423448145 |
| 11. | CASP9 | BART12, BART17-5p, BART19-3p, BART21-5p, BART3, BART4, BART6-3p, BART7* | -3.377407397 |
| 12. | BID | BART1-5p, BART16, BART17-3p, BART18-3p, BART19-3p, BART20-3p, BART3*, BART4, BART6-3p, BHRF1-2, BHRF1-3 | -3.276606757 |
| 13. | IL1B | BART14*, BART15, BART2-5p, BART6-3p | -3.276606756 |
| 14. | AKT1 | BART11-5p, BART12, BART1-5p, BART17-3p, BART5, BART7, BHRF1-3, hbv-miR-B14RC-3p | -3.27586137 |
| 15. | PRKAR2B | BART10, BART13, BART14, BART15, BART1-5p, BART16, BART19-3p, BART22, BART3*, BART6-3p, BART7, BHRF1-1 | -3.239059262 |
| 16. | BCL2L1 | BART10*, BART11-3p, BART12, BART13*, BART13, BART1-5p, BART20-5p, BART3, BART4, BART5*, BART6-3p, BART7*, BART8*, BHRF1-2* | -2.971486949 |
| 17. | PIK3R2 | BART12, BART13*, BART1-5p, BART16, BART17-5p, BART18-3p, BART6-3p, BART7, BHRF1-1 | -2.965520221 |
| 18. | ENDOG | BART10*, BART1-3p, BART19-5p | -2.891316601 |
| 19. | IKBKG | BART11-5p, BART17-5p, BART5 | -2.790417741 |
| 20. | BAX | BART22, BART2-3p | -2.742663852 |
| 21. | TRAF2 | BART10*, BART17-3p, BART18-5p, BART4*, BART4, BART5*, BART6-3p | -2.720825281 |
| 22. | TP53 | BART10*, BART11-3p, BART11-5p, BART12, BART14, BART15, BART1-5p, BART18-5p, BART21-3p, BART21-5p, BART2-5p, BART5*, BART6-5p, BART7, BART8* | -2.70715363 |
| 23. | IRAK1 | BART10*, BART11-5p, BART12, BART13*, BART17-5p, BART3, BART4, BART5*, BART6-3p, BHRF1-1 | -2.633412779 |
| 24. | FADD | BART10*, BART12, BART13, BART1-5p, BART17-5p, BART3*, BART7*, BART7, BHRF1-2*, | -2.610709565 |
| 25. | CYCS | BART10*, BART10, BART12, BART1-3p, BART14, BART15, BART1-5p, BART16, BART17-3p, BART17-5p, BART18-5p, BART19-3p, BART19-5p, BART20-5p, BART22, BART2-5p, BART3, BART4*, BART4, BART5, BART6-5p, BART9*, BART9, BHRF1-1, BHRF1-2*, BHRF1-2 | -2.560022662 |
| 26. | AKT2 | BART10*, BART11-3p, BART12, BART13*, BART1-3p, BART14, BART15, BART16, BART17-5p, BART20-3p, BART20-5p, BART2-3p, BART3, BART4*, BART5*, BART5, BART6-3p, BART7, BART9*, BHRF1-1, BHRF1-3 | -2.516024057 |
| 27. | DFFB | BART10, BART11-3p, BART11-5p, BART13, BART1-3p, BART14, BART17-5p, BART18-3p, BART20-5p, BART22, BART2-3p, BART3, BART4, BART6-5p, BART7, BART9*, BART9, BHRF1-1 | -2.409057053 |
| 28. | PRKAR2A | BART11-5p, BART12, BART13, BART14, BART18-5p, BART2-5p, BART4 | -2.321000876 |
| 29. | DFFA | BART10*, BART10, BART11-3p, BART12, BART1-3p, BART14, BART1-5p, BART16, BART17-3p, BART18-3p, BART18-5p, BART19-3p, BART20-5p, BART4, BART5*, BHRF1-1 | -2.187673028 |
| 30. | IRAK2 | BART11-5p, BART14, BART17-3p, BART19-3p, BART19-5p, BART22, BART9, BHRF1-1 | -2.083961677 |
| 31. | PRKAR1A | BART10*, BART10, BART11-5p, BART14*, BART14, BART16, BART17-3p, BART18-3p, BART19-3p, BART21-3p, BART22, BART2-5p, BART4*, BART5, BART6-3p, BART6-5p, BART8*, BART9*, BART9, BHRF1-2*, BHRF1-2, BHRF1-3 | -2.06536446 |
| 32. | IKBKB | BART12, BART13, BART14, BART15, BART16, BART17-5p, BART20-5p, BART21-5p, BART22, BART2-3p, BART2-5p, BART3*, BART4*, BART6-3p, BART8* | -1.950163181 |
| 33. | NFKB1 | BART1-5p, BART20-5p, BART3*, BART5* | -1.598534851 |
| 34. | BNIP3L | BART13, BART14, BART15, BART16, BART17-3p, BART18-5p, BART19-3p, BART19-5p, BART21-3p, BART22, BART2-5p, BART3, BART4*, BART4, BART6-5p, BART7, BART8*, BART9*, BART9, BHRF1-1, BHRF1-2, BHRF1-3 | -1.52453427 |
| 35. | PPP3CB | BART10*, BART14, BART18-3p, BART18-5p, BART20-5p, BART2-3p, BART4*, BART7, BART8*, BART9, BHRF1-2*, BHRF1-2, BHRF1-3 | -1.497605943 |
| 36. | MYD88 | BART10*, BART11-5p, BART12, BART13*, BART13, BART14, BART15, BART1-5p, BART16, BART17-3p, BART17-5p, BART19-3p, BART19-5p, BART22, BART2-3p, BART2-5p, BART5*, BART5, BART6-3p, BART6-5p, BART8*, BHRF1-2 | -1.446963767 |
| 37. | PIK3CD | BART11-3p, BART11-5p, BART12, BART1-3p, BART19-3p, BART19-5p, BART20-3p, BART21-3p, BART2-5p, BART3, BART4, BART5*, BART5, BART6-3p, BART9 | -1.440040431 |
| 38. | BNIP3 | BART15, BART17-5p, BART19-3p, BART6-5p | -1.276606757 |
| 39. | TNFRSF10A | BART17-5p, BHRF1-1 | -1.222386366 |
| 40. | MAP3K14 | BART13, BART19-3p, BART22, BART4, BART5*, BHRF1-1 | -1.132216847 |
| 41. | PRKX | BART10*, BART10, BART11-3p, BART11-5p, BART14, BART15, BART16, BART17-3p, BART18-5p, BART19-3p, BART19-5p, BART20-5p, BART22, BART2-3p, BART2-5p, BART3, BART4*, BART4, BART5*, BART5, BART6-5p, BART7*, BART7, BART8*, BHRF1-2*, BHRF1-2 | -1.011417466 |
| 42. | TNFRSF1A | BART1-3p, BART5 | -0.942869359 |
| 43. | PPP3R1 | BART11-5p, BART12, BART1-3p, BART14, BART16, BART19-3p, BART19-5p, BART20-3p, BART21-3p, BART22, BART2-5p, BART3*, BART8*, BART9, BHRF1-2 | -0.890999977 |
| 44. | NFKBIA | BART13, BART4*, BART5*, BART5, BART8*, BART9 | -0.86627626 |
| 45. | CSF2RB | BART10*, BART11-5p, BART12, BART1-3p, BART17-3p, BART18-3p, BART18-5p, BART19-3p, BART20-5p, BART21-3p, BART22, BART2-5p, BART4*, BART4, BART5*, BART7*, BART7, BART9, BHRF1-1, BHRF1-3 | -0.735515754 |
| 46. | RIPK1 | BART10, BART11-3p, BART11-5p, BART12, BART13, BART14, BART1-5p, BART17-3p, BART17-5p, BART19-3p, BART20-3p, BART2-3p, BART6-3p, BART6-5p, BART8*, BHRF1-2* | -0.709566164 |
| 47. | PIK3R5 | BART13*, BART17-3p, BART17-5p, BART18-5p, BART20-3p, BART20-5p, BART3, BART6-3p, BART6-5p, BHRF1-1 | -0.691644255 |
| 48. | CASP7 | BART12, BART15, BART19-3p, BART22, BART5, BART6-3p, BART8*, BART9* | -0.668924179 |
| 49. | CHUK | BART15, BART1-5p, BART22, BART7 | -0.655985124 |
